# Zoliflodacin and gepotidacin cross-resistance in *Neisseria gonorrhoeae*

**DOI:** 10.1101/2025.11.25.690462

**Authors:** Aditi Mukherjee, Sofia OP Blomqvist, David Helekal, Apabrita A Das, Samantha G Palace, Yonatan H Grad

**Affiliations:** Department of Immunology and Infectious Diseases, Harvard T. H. Chan School of Public Health, Boston, Massachusetts, USA; Cardiovascular Medicine Division, Brigham and Women’s Hospital, Boston, Massachusetts, USA; Harvard Medical School, Boston, Massachusetts, USA

## Abstract

Resistance to zoliflodacin, a first-in-class antibiotic for gonorrhea treatment, can occur through *gyrB*^D429N^, but this mutation’s impact on fitness and resistance to other topoisomerase targeting drugs, including gepotidacin, have been unclear. Here, we show that *gyrB*^D429N^ confers cross-resistance to gepotidacin in some clinical isolates and that its fitness effect varies with strain background. These findings inform strategies for introducing the new topoisomerase inhibitors into clinical use and for surveillance of resistance.

---

The World Health Organization estimated over 82 million new *Neisseria gonorrhoeae* infections in 2020, with the highest burden of antimicrobial resistance reported in Asia^1^. The prevalence of ciprofloxacin resistance in Asia is at or near 100%^2^, and ceftriaxone resistance has reached 30% in some regions^3^. Two new, first-in-class antibiotics, zoliflodacin and gepotidacin, successfully completed phase 3 trials for uncomplicated gonorrhea^4, 5^. Fluoroquinolones and gepotidacin target the A subunit of DNA gyrase (*gyrA*) and A subunit of topoisomerase IV (*parC*)^6^, and zoliflodacin targets the *gyrB* subunit of DNA gyrase^7^. Resistance to ciprofloxacin arises from mutations in codons 91 and 95 in *gyrA* and 86-91 in *parC*^8^. Resistance to zoliflodacin has been linked to substitutions in *gyrB*, including D429N, K450N, and K450T^9, 10^. Additionally, the S467N mutation in *gyrB* appears to act as a stepping-stone mutation to zoliflodacin resistance but does not itself confer resistance to zoliflodacin^11^. In the phase 2 trial for gepotidacin treatment of gonorrhea, gepotidacin resistance arose via spontaneous *gyrA*^A92T^ mutation in two isolates that harbored the allele *parC*^D86N^, which both contributes to ciprofloxacin resistance and functions as a stepping stone to gepotidacin resistance^12-15^. Zoliflodacin and gepotidacin are active against ciprofloxacin-resistant *N. gonorrhoeae* and cross-resistance among these three topoisomerase-targeting drugs has not been reported^15, 16^. However, the overlapping targets raise concern that existing genetic diversity in topoisomerase components among prevalent ciprofloxacin-resistant lineages may increase the likelihood of acquiring resistance to zoliflodacin and gepotidacin, either by enabling cross-resistance or by ameliorating resistance-associated fitness costs.

To determine the effects of *gyrB*^D429N^ on antimicrobial susceptibility across genetic backgrounds, we introduced this mutation into each of 9 *N. gonorrhoeae* clinical isolates, representing common lineages (**Supplementary Figure 1, Supplementary Table 1**). These were selected to capture the genetic diversity with respect to alleles that define ciprofloxacin-resistance genotypes such as *gyrA*^91F, 95G/A^ and *parC* allelic variation at codon positions 86, 87, and 91. In each isolate, *gyrB*^D429N^ increased zoliflodacin MICs 16- to 32-fold, confirming the ability of this substitution to confer zoliflodacin resistance independent of background (**Figure 1A**).

**Figure 1:**
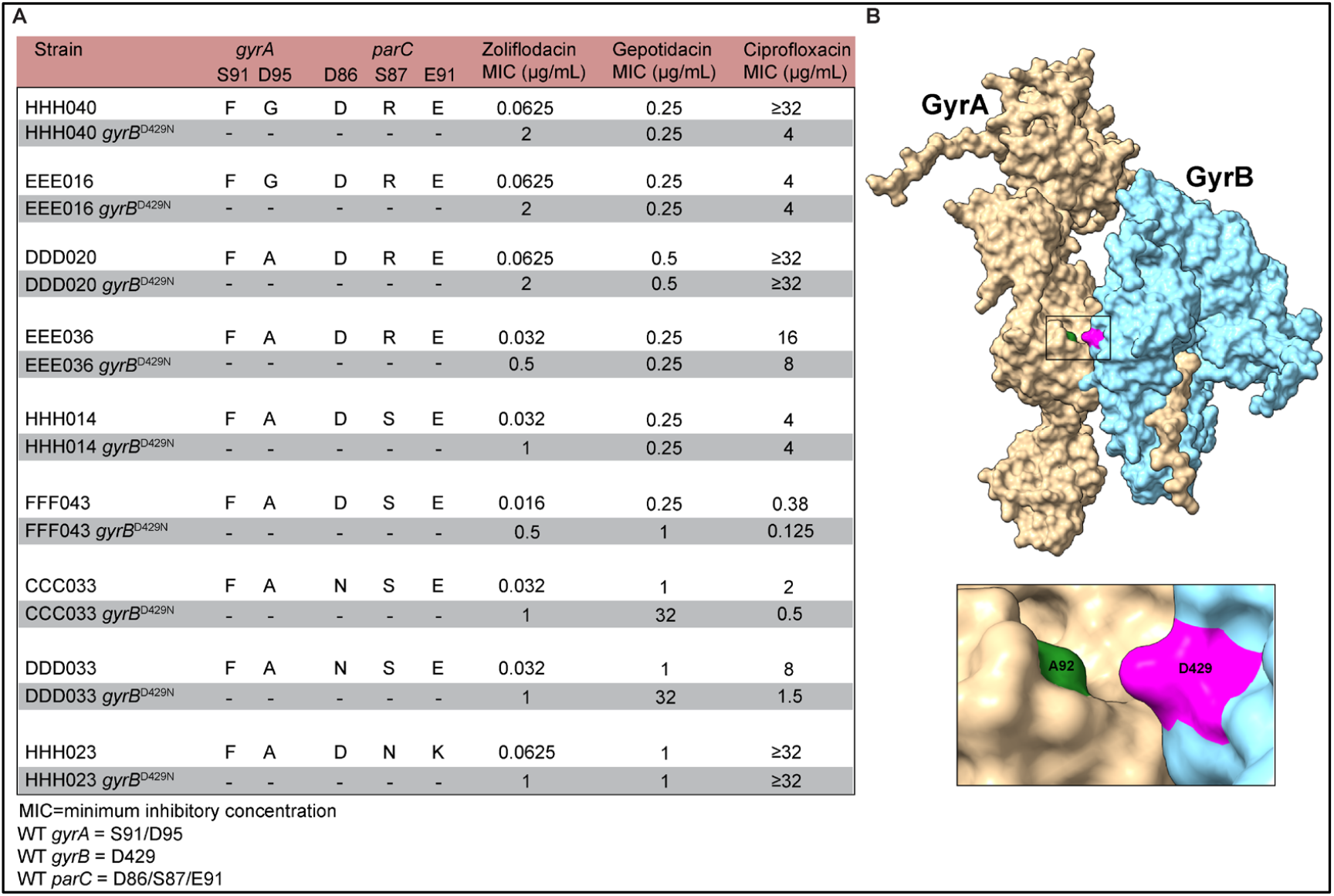
**(A)** Zoliflodacin, gepotidacin, and ciprofloxacin MICs of *N. gonorrhoeae* clinical isolates and their *gyrB*^D429N^ variants. **(B)** Top: ColabFold predicted structure of GyrA (light brown) and GyrB (light blue) docked into a heterodimeric complex. Bottom: Magnified view of the GyrA-GyrB interface highlighting residues GyrA^A92^ (green) and GyrB^D429^ (magenta).

In 6 out of 9 strains, *gyrB*^D429N^ did not affect gepotidacin resistance, as expected. However, *gyrB*^D429N^ conferred large increases in gepotidacin resistance in the remaining three isolates: a 32-fold MIC increase in two isolates (CCC033 and DDD033) and a 4-fold MIC increase in the third (FFF043). Both CCC033 and DDD033 harbor the *parC*^D86N^ mutation that contributes to gepotidacin resistance when in combination with *gyrA*^A92T^, but Sanger sequencing confirmed that the zoliflodacin- and gepotidacin-resistant *gyrB*^D429N^ derivatives of these strains did not spontaneously acquire the *gyrA*^A92T^ mutation. FFF043 lacks variants in the *parC* sites associated with quinolone resistance, indicating other sites may also contribute to zoliflodacin/gepotidacin cross-resistance (**Figure 1A**).

To understand the mechanism by which *gyrB*^D429N^ can increase gepotidacin MICs, we created a structural model of the *N. gonorrhoeae* GyrA-GyrB heterodimer. The predicted complex (**Supplementary Figure 2**) showed that GyrA residues 91, 92, and 95 associated with resistance to ciprofloxacin and gepotidacin cluster along the same surface. On the GyrB subunit, zoliflodacin resistance-associated residues 429, 450, and 467 are positioned along a contiguous region. GyrA^92^, the position at which gepotidacin resistance mutations were observed following treatment failure in the Phase 2 clinical trial, lies in close spatial proximity to GyrB^429^ (**Figure 1B**). This spatial arrangement suggests that alterations in *gyrB*^D429^ may alter gepotidacin binding via a similar mechanism to the known resistance mutation *gyrA*^A92T^. High-level gepotidacin resistance from *gyrA*^A92T^ only emerged in *N. gonorrhoeae* strains that also harbored the *parC*^D86N^ mutation, similar to the background-dependent pattern of *gyrB*^D429N^ cross-resistance to gepotidacin that we observed. Taken together, these results point to a working model in which gepotidacin resistance requires target site mutations at both known drug targets: the *parC*^D86N^ mutation to preserve topoisomerase IV function, and either the *gyrA*^A92T^ or the *gyrB*^D429N^ mutation to preserve gyrase function.

The effects of *gyrB*^D429N^ on ciprofloxacin susceptibility were variable. Five isolates (EEE016, DDD020, EEE036, HHH014 and HHH023) maintained ciprofloxacin resistance in the presence of *gyrB*^D429N.^ In the remaining four isolates, ciprofloxacin MICs were reduced by at least 2-fold and in one case as much as ≥8-fold (HHH040) (**Figure 1A**). The differential effect of *gyrB*^D429N^ on ciprofloxacin MICs does not appear to be explained by *gyrA* or *parC* genotypes.

To evaluate the fitness impact of *gyrB*^D429N^, we evaluated *in vitro* growth of each strain and its *gyrB*^D429N^ derivative in antibiotic-free gonococcal medium in monoculture (**Supplementary Figure 3**,**4**) and in competition assays (**Figure 2; Supplementary Figure 5**). Introduction of *gyrB*^D429N^ imposed a fitness cost in most of the isolates (**Figure 2, Supplementary Figure 5**). The strongest fitness costs were observed in HHH040 and EEE016, both of which carry *gyrA*^91F, 95G^ and *parC*^S87R^. In contrast, the introduction of the *gyrB*^D429N^ mutation into EEE036 resulted in a fitness advantage (**Figure 2D; Supplementary Figure 5D**). Whole-genome sequencing confirmed that no off-target mutations occurred in the EEE036 *gyrB*^D429N^ strain (**Supplementary Table 2**). EEE036 carries the same combination of *gyrA*^91F/95A^ and *parC*^S87R^ alleles found in DDD020, in which the *gyrB*^D429N^ mutation incurred a substantial fitness cost, indicating that EEE036 harbors another genetic variant that accommodates the *gyrB*^D429N^ mutation.

**Figure 2:**
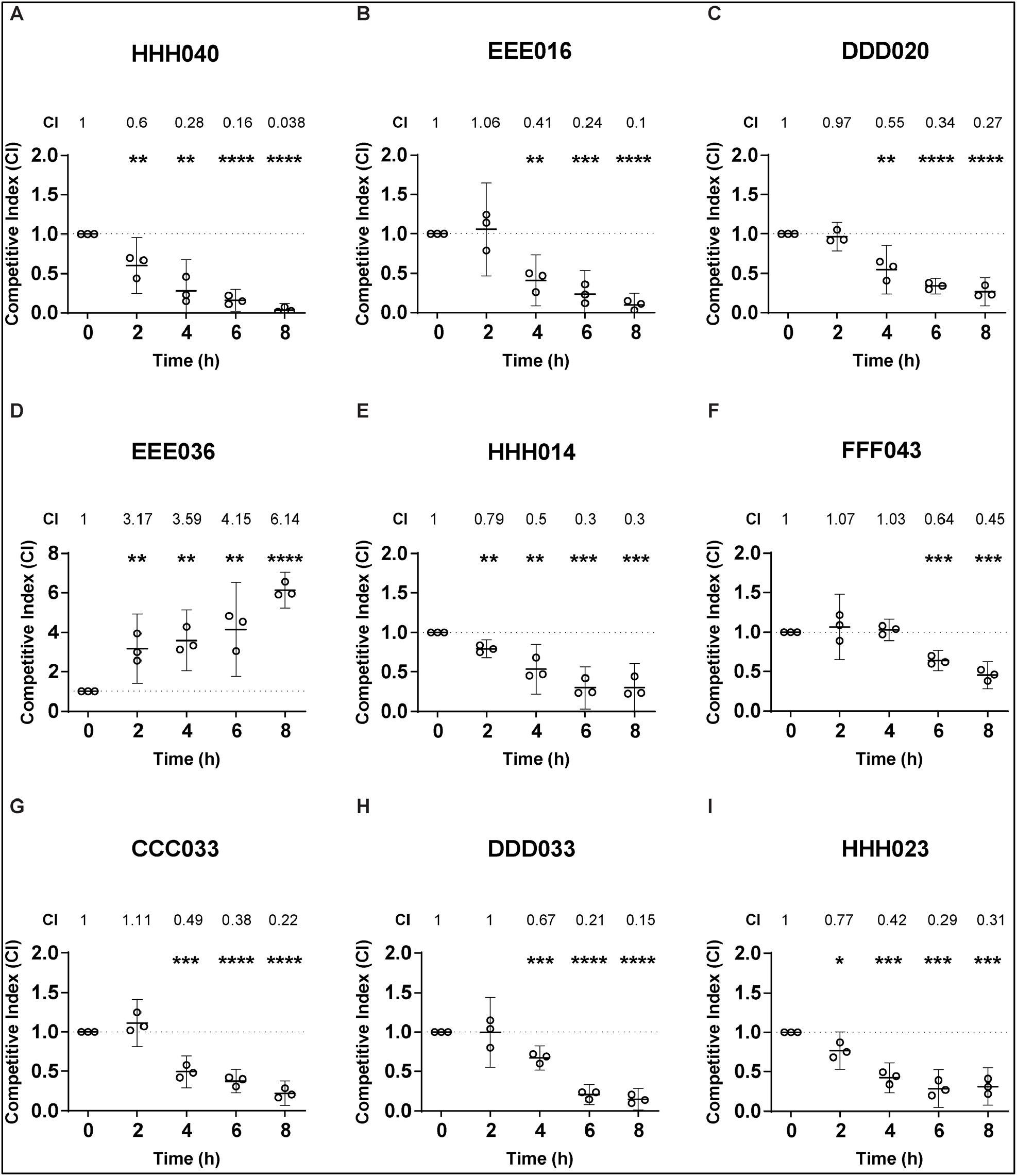
Relative fitness of the *gyrB*^D429N^ mutant in 9 clinical isolate backgrounds. Y-axes show the Competitive Index (CI) of each *gyrB*^D429N^ mutant relative to its parental strain during competitive growth *in vitro*. In all cases, the *gyrB*^D429N^ mutant carried a kanamycin marker and was cocultured with its unmarked parental strain. (A) HHH040: p = 0.009, 0.002, <0.0001, <0.0001 respectively for 2, 4, 6 and 8 hours. (B) EEE016: p = 0.7, 0.001, 0.0004, <0.0001 respectively for 2, 4, 6 and 8 hours. (C) DDD020: p = 0.47, 0.003, <0.0001, <0.0001 respectively for 2, 4, 6 and 8 hours. (D) EEE036: p = 0.006, 0.002, 0.005, <0.0001 respectively for 2, 4, 6 and 8 hours. (E) HHH014: p = 0.001, 0.003, 0.0003, 0.0006, respectively for 2, 4, 6 and 8 hours. (F) FFF043: p = 0.53, 0.4, 0.0003, 0.0002, respectively for 2, 4, 6 and 8 hours. (G) CCC033: p = 0.18, 0.0004, <0.0001, <0.0001, respectively for 2, 4, 6 and 8 hours. (H) DDD033: p = 0.98, 0.0008, <0.0001, <0.0001, respectively for 2, 4, 6 and 8 hours. (I) HHH023: p = 0.01, 0.002, 0.0002, 0.0002, respectively for 2, 4, 6 and 8 hours. n = 3, representative of three independent experiments performed in absence of any antibiotic pressure. Error bars represent mean with 95% confidence interval. Statistically significant differences in competitive indices compared to time 0 were analyzed using an unpaired Student’s *t-test*, indicated *p ≤ 0.05, **p ≤ 0.01, ***p ≤ 0.001 and ****p ≤ 0.0001.

The zoliflodacin-resistance mutation *gyrB*^D429N^ invariably increased zoliflodacin MIC, but modulated susceptibility to ciprofloxacin and gepotidacin in a strain-dependent manner. While resistance to gepotidacin has been observed through a combination of *parC*^D86N^ and *gyrA*^A92T13-15^, our study suggests that *gyrB*^D429N^ can also yield gepotidacin resistance (MIC=32 µg/mL) in the *parC*^D86N^ background. This raises concern for the emergence of cross-resistant strains, given that regions where zoliflodacin may be introduced first due to ceftriaxone resistance also have reported a high prevalence of *parC*^D86N^: 18.9% in Vietnam^17^, 36.7% in Cambodia^18^, and 46.5% in Thailand^19^. Cross-resistance would be expected to shorten the duration of the new drugs’ clinically useful lifespans^20, 21^.

The fitness consequences of resistance mutations play a major role in determining how quickly resistance will arise and spread. We found that the fitness effects of *gyrB*^D429N^ varied with genetic background, ranging from deleterious to advantageous. While *gyrB*^D429N^ imposed a fitness cost in most clinical isolates we tested, the fitness advantage it conferred in isolate EEE036 demonstrates that this resistance mutation is not universally costly. The translation of *in vitro* fitness to fitness within the context of human infections is uncertain, although other studies have shown a correlation^22, 23^. If this unexpected fitness benefit of *gyrB*^D429N^ translates to *in vivo* settings, this observation indicates that some zoliflodacin-resistant strains could persist and spread even in the absence of drug pressure and further suggests that resistance could be similarly stabilized in other lineages by the evolution of compensatory mutations.

Our results have several additional implications. First, the susceptibility and fitness effects of resistance mutations are modulated by genetic background, indicating that additional factors remain to be discovered. Second, standard surveillance approaches that track only individual loci (e.g., *gyrB*) will likely be insufficient for monitoring resistance phenotypes. Third, early recognition of background-dependent resistance emergence could allow for regionally tailored treatment guidelines, preventing widespread therapeutic failure^24^.

Finally, while early *in vitro* and clinical experiences with the two new drugs identified some resistance mutations, our findings indicate that other, in this case cross-resistance conferring, mutations may arise. More comprehensive analysis and vigilant surveillance for phenotypic resistance as the new drugs are rolled out will be important to inform optimal clinical use and to maximize their public health benefit.

## Methods

### Phylogenetic analysis

We assembled a dataset of published genomes ^19, 25-27^ and isolates selected from those collected by the Centers for Disease Control and Prevention’s (CDC) 2024 surveillance efforts.

To obtain the genomes of the CDC isolates, we searched the NCBI Pathogen Detection database (https://www.ncbi.nlm.nih.gov/pathogens/) for *Neisseria gonorrhoeae* isolate genome sequences collected by the US CDC with a collection date between January 1, 2024, and December 31^st^, 2024 (“Home - pathogen detection - NCBI”). We filtered out isolates that were not identified as a GCWGS isolate, the designation given to isolates collected in the context of CDC *N. gonorrhoeae* surveillance. The accession numbers of the publicly available isolates used for the phylogenetic tree are included in **Supplementary Table 1**.

Reference-based mapping to NCCP11945 (NC_011035.1) was done using BWA-MEM v0.7.17^28^. We used Pilon v1.23 to call variants (minimum mapping quality: 20, minimum coverage: 10X)^29^ after marking duplicate reads with Picard v2.20.1 (https://broadinstitute.github.io/picard/) and sorting reads with SAMtools v1.17^30^. We generated pseudogenomes by incorporating variants supported by at least 90% of reads and sites with ambiguous alleles into the reference genome sequence. The pseudogenomes were then used as the input alignment for subsequent phylogenetic reconstruction.

We used GUBBINS v3.4.3^31^ to estimate recombining regions and IQTREE v2.4.0^32^ for phylogenetic reconstruction. We used MODELFINDER^33^ to select an optimal molecular clock model. The phylogenetic tree was visualised using iTOL v7^34^.

We generated pseudogenomes and used these to call *parC*^86^ variants as described above for isolates published previously^17, 18^.

### Molecular docking

Protein sequences of *N. gonorrhoeae* GyrA and GyrB were retrieved from UniProt (accessions P48371 and P22118, respectively). Tertiary structures of GyrA and GyrB monomers were predicted independently using ColabFold v1.5^35^ with AlphaFold2-multimer presets and default parameters. For each monomer, the top-ranked model was selected from ColabFold results. GyrA and GyrB monomers were docked with ClusPro 2.0 protein-protein docking server^36^ using balanced scoring model and the highest ranked structure was used for structural analysis. ChimeraX v1.10.1^37^ was used for visualization of protein interfaces.

### *N. gonorrhoeae* culture conditions

*N. gonorrhoeae* was cultured on GCB agar (Difco) supplemented with Kellogg’s supplement (GCB-K) at 37ºC with 5% CO_2_^38^. Growth curve and pairwise competition experiments were conducted in liquid GCP medium containing 15 g/L proteose peptone 3 (Thermo Fisher), 1 g/L soluble starch, 1 g/L KH_2_PO_4_, 4 g/L K_2_HPO_4_, and 5 g/L NaCl (Sigma-Aldrich) with Kellogg’s supplement^39^ with agitation at 37ºC with 5% CO_2_.

### Generation of isogenic *N. gonorrhoeae gyrB* mutant strains

Strains, plasmids and primers used in this study are listed in **Supplementary Table 1**.

The mutant *gyrB*^D429N^ allele was amplified using primers AM_1 (F) and AM_2 (R) from the genomic DNA of a previously reported experimentally evolved *N. gonorrhoeae* GCGS0481 strain that acquired the *gyrB*^D429N^ mutation under ciprofloxacin pressure^9^ and introduced into the selected isolates by electroporation^22^. Individual colonies were selected on GCB-K plates from within the zone of inhibition created by a dried droplet of 4 μg/mL zoliflodacin. Transformants were verified by Sanger sequencing of the *gyrA* and *gyrB* loci and further examined by whole genome sequencing. Genomic DNA from parent and mutant strains were purified using an Invitrogen PureLink Genomic DNA Mini Kit (K182001), prepared for sequencing using Oxford Nanopore Technologies Native Barcoding Kit 24 V14 (SQK-NBD114.24), and sequenced on an Oxford Nanopore Technologies R10.4.1 flow cell followed by basecalling with Dorado v0.8.1 (https://github.com/na-noporetech/dorado/tree/release-v0.8) with super accuracy. Genome assemblies were created with Autocycler v0.2.1^40^ with 4 read subsets at 25x minimum depth and using the assemblers Canu v2.2^41^, Flye v2.9.5^42^, Miniasm v0.3^43^, NECAT v0.0.1^44^, NextDenovo v2.5.2^45^, and Raven v1.8.3^46^. For each pair of strains, *gyrB*^429N^ reads were mapped to the *gyrB*^429D^ parent’s de novo assembly with SAMtools v1.21^30^ and Minimap2 v2.28^47^. Variants between *gyrB*^429D^ and *gyrB*^429N^ strains were identified using Pilon v1.24^29^ with a minimum alignment mapping quality of 20 and minimum depth of 10. The resulting VCF was filtered to keep only variants with an allele frequency greater than 0.9 and a depth greater than 5. Assemblies were annotated with Prokka v1.14.6^48^. Basecalled reads were uploaded to the NCBI SRA and are available at PRJNA1368854. All variants detected in *gyrB*^D429N^ transformants compared to their parent strains are summarized in **Supplementary Table 2**.

### Antibiotic susceptibility testing

Antibiotic susceptibility testing was performed on GCB-K agar via Etest (BioMerieux) for ciprofloxacin or agar dilution for zoliflodacin and gepotidacin. All MIC results reported are the mean of three independent experiments.

### Measurement of growth and competitive fitness of *gyrB* variants

To measure the growth of each strain and its *gyrB*^D429N^ derivative, overnight cultures on GCB-K plates were diluted to an optical density (OD) of 0.1 (600nm) and grown in antibiotic-free GCP medium with Kellogg’s supplement for 8 hours. OD_600_ of each strain was measured at 2, 4, 6 and 8 hours. At each timepoint, cultures were also serially diluted and plated on GCB-K agar plates to measure colony forming units (CFUs) as a proxy for viable bacteria.

For competition assays, kanamycin resistance was introduced into each strain in the panel of clinical isolates and each *gyrB*^D429N^ derivative by electroporation with pDR53, which integrates an *aphA3* marker onto the chromosome between *lctP* and *aspC*^22^. Transformants were selected on GCB-K agar supplemented with 70 µg/ml kanamycin. In pairwise competition experiments, paired strains (one kanamycin-sensitive and one kanamycin-resistant strain) were mixed at a ratio of 1:1 by OD_600_, diluted to OD_600_ ∼0.1, and co-cultured in antibiotic-free GCP media with Kellogg’s supplement for 8 hours. At each timepoint, cultures were serially diluted and plated on both GCB-K agar and GCB-K agar supplemented with 70 µg/ml kanamycin. Plates were incubated overnight at 37°C 5% CO_2_. Colonies on each plate were quantified, and the competitive index was calculated at each timepoint as (*R*_*t*_ / *S*_*t*_) / (*R*_*0*_ / *S*_*0*_), where *R*_*t*_ and *S*_*t*_ are the proportions of kanamycin-resistant and kanamycin-sensitive strains, respectively, at time *t* and *R*_*0*_ and *S*_*0*_ are the proportions of kanamycin-resistant and kanamycin-sensitive strains at time 0. A competitive index value of 1 indicates equal fitness between strains, >1 indicates the kanamycin-resistant mutant is more fit than the parental strain, and <1 indicates the mutant is less fit. Statistical analysis of competitive index measurements was performed using an unpaired two-sided Student’s *t-test*. All competition experiments were performed by competing unmanipulated, kanamycin susceptible clinical isolates against kanamycin-marked versions of their isogenic *gyrB*^D429N^ derivatives (**Figure 2**) as well as by competing kanamycin-marked parental strains against kanamycin susceptible *gyrB*^D429N^ strains (**Supplementary Figure 5**) to ensure that the kanamycin marker did not contribute to the fitness differences reported here.

## Acknowledgements

We would like to thank members of the Grad lab for helpful discussions and feedback on this manuscript. This work was supported by NIH R01 AI132606 and R01 AI153521 grants to Y.H.G.

## Author contributions

AM, SGP and YHG conceptualized the study. AM, AAD, SOPB and DH acquired/analyzed the data. AM, SGP and YHG wrote the original draft, and all the authors reviewed and edited the manuscript.

## Supplementary Files

**Supplementary Figure 1:**
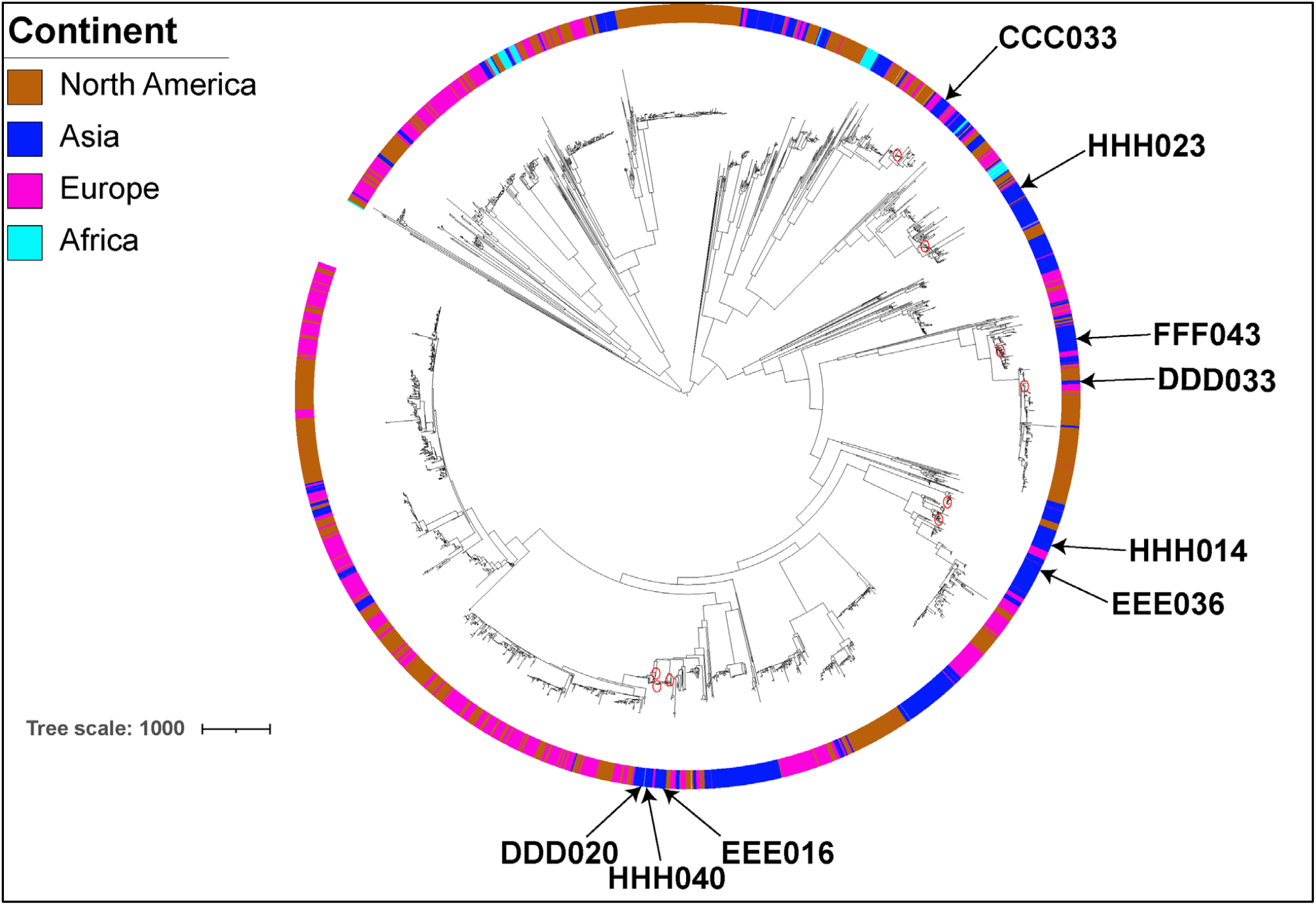
Phylogenetic tree of *N. gonorrhoeae* clinical isolates. Tree scale represents the recombination-adjusted number of single-nucleotide polymorphisms. Isolates used in this study are marked by arrows. The outer ring represents continents (North America, Asia, Europe, Africa) of the origin of the isolates.

**Supplementary Figure 2:**
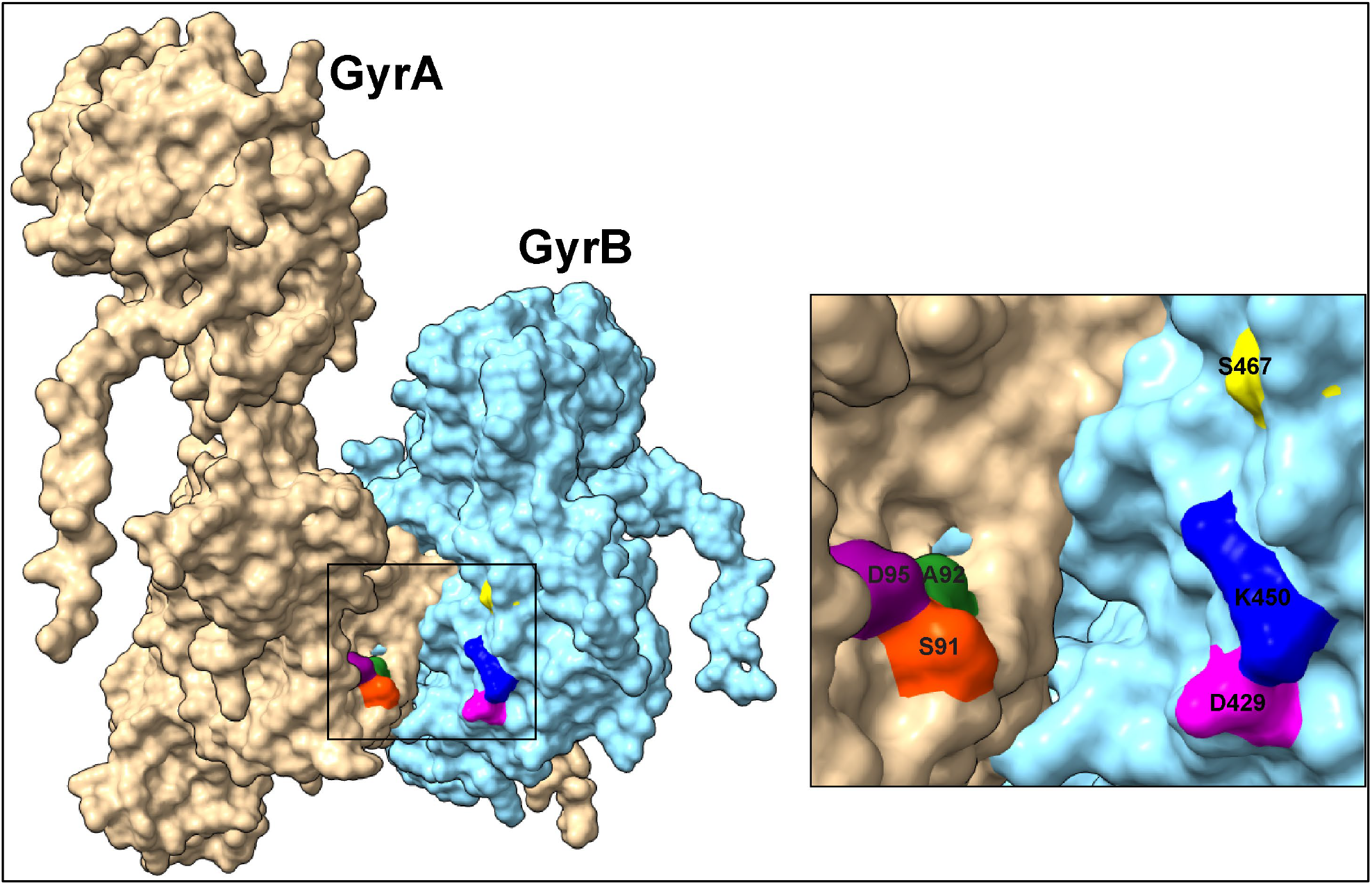
Left: ColabFold predicted structure of wildtype GyrA (light brown) and wildtype GyrB (light blue) docked into a heterodimeric complex. Right: Magnified view of the GyrA-GyrB interface highlighting residues GyrA^S91^ (orange), GyrA^A92^ (green), GyrA^D95^ (purple) and GyrB^D429^ (magenta), GyrB^K450^ (blue), GyrB^S467^ (yellow).

**Supplementary Figure 3:**
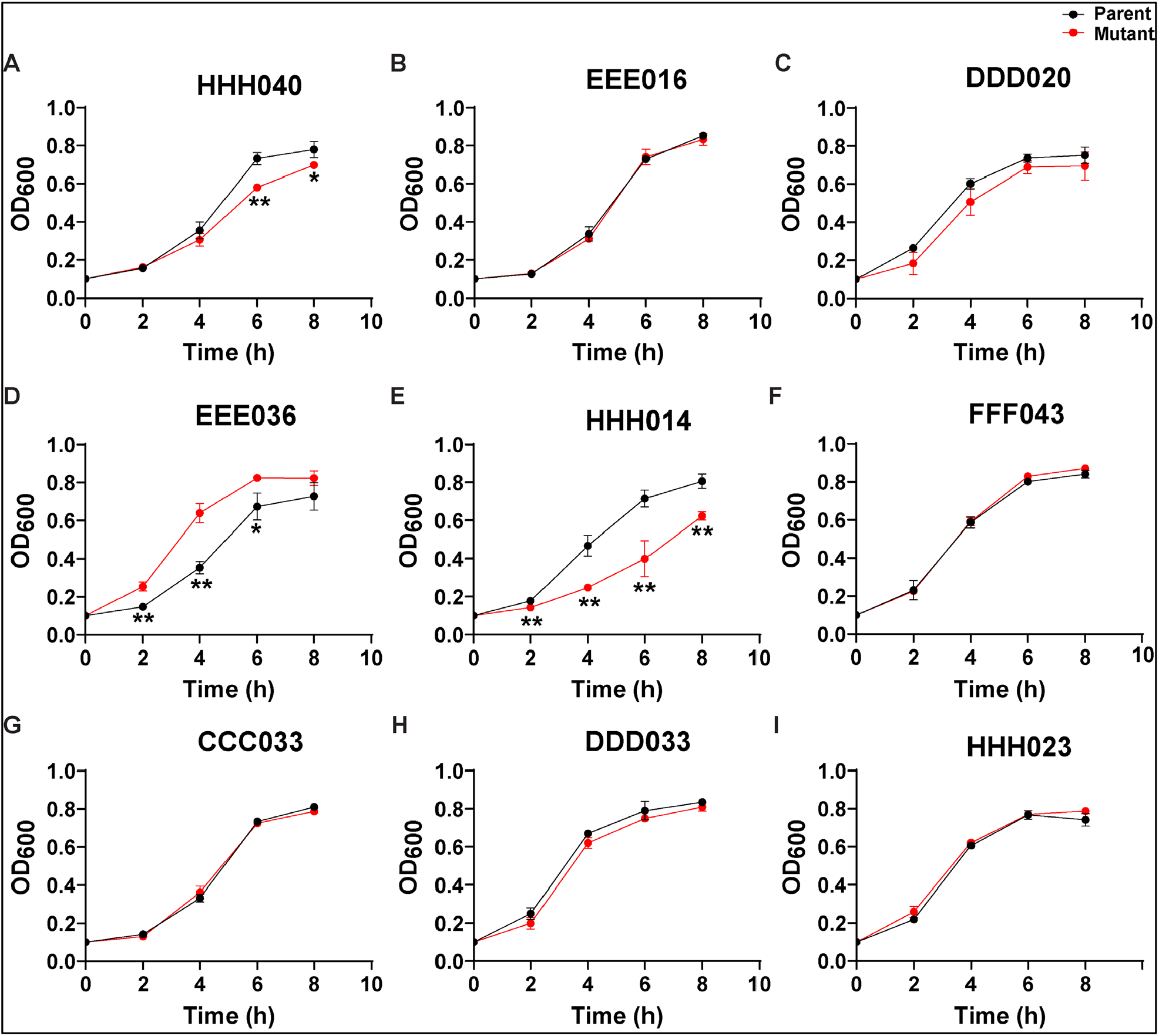
*In vitro* growth kinetics of isolates with and without *gyrB*^D429N^, measured by spectrophotometry. All isolates (black) and their isogenic *gyrB*^D429N^ mutants (red) were cultured separately in liquid GCP media supplemented with Kellogg’s supplement at a starting absorbance reading at 600nm (OD_600_) = 0.1. OD_600_ readings at 0, 2, 4, 6 and 8 hours timepoint are plotted. (A) HHH040 and HHH040 *gyrB*^429N^. p = 0.44, 0.2, 0.002, 0.03, respectively for 2, 4, 6 and 8 hours. (B) EEE016 and EEE016 *gyrB*^429N^. (C) DDD020 and DDD020 *gyrB*^429N^. (D) EEE036 and EEE036 *gyrB*^429N^. p = 0.002, 0.001, 0.02, 0.11, respectively for 2, 4, 6 and 8 hours. (E) HHH014 and HHH014 *gyrB*^429N^. p = 0.005, 0.003, 0.006, 0.002, respectively for 2, 4, 6 and 8 hours. (F) FFF043 and FFF043 *gyrB*^429N^. (G) CCC033 and CCC033 *gyrB*^429N^. (H) DDD033 and DDD033 *gyrB*^429N^. (I) HHH023 and HHH023 *gyrB*^429N^. n = 3, representative of three independent experiments. Error bars represent SDs between three biological replicates. Statistical significance was determined by unpaired two-sided Student’s *t-test* and are indicated *p ≤ 0.05 and **p ≤ 0.01.

**Supplementary Figure 4:**
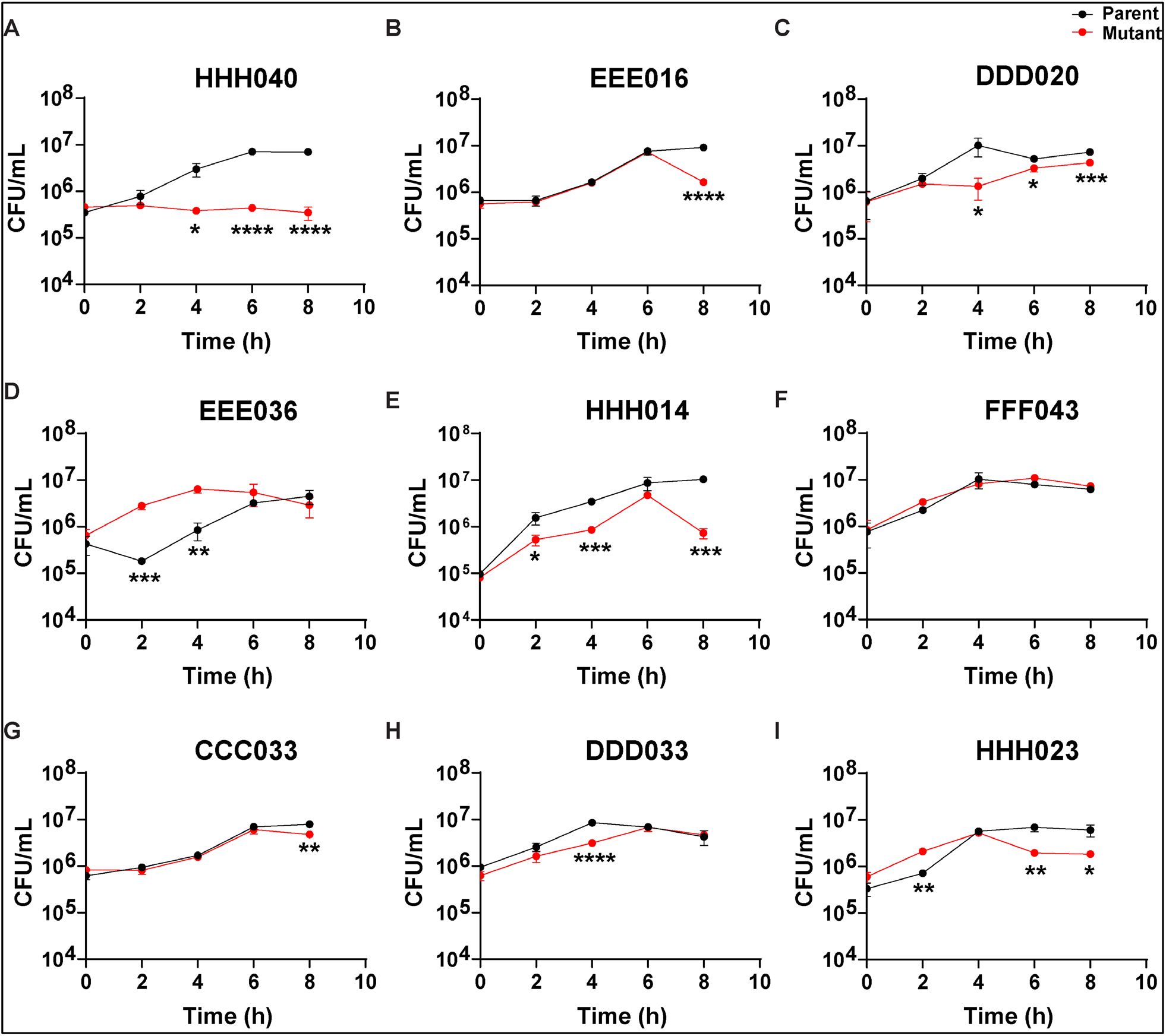
*In vitro* growth kinetics of isolates with and without *gyrB*^D429N^, measured by dilution plating. All isolates (black) and their isogenic *gyrB*^D429N^ mutants (red) were cultured separately in liquid GCP media supplemented with Kellogg’s supplement at a starting absorbance reading at 600nm (A_600_) = 0.1. Dilutions were plated on GCB-K plates at 0, 2, 4, 6 and 8 hours, colonies were counted after overnight growth, and CFUs were calculated. (A) HHH040 and HHH040 *gyrB*^429N^. p = 0.14, 0.01, <0.0001, <0.0001, respectively for 2, 4, 6 and 8 hours. (B) EEE016 and EEE016 *gyrB*^429N^. p = 0.7, 0.76, 0.73, <0.0001, respectively for 2, 4, 6 and 8 hours. (C) DDD020 and DDD020 *gyrB*^429N^. p = 0.23, 0.03, 0.01, 0.001, respectively for 2, 4, 6 and 8 hours. (D) EEE036 and EEE036 *gyrB*^429N^. p = 0.0009, 0.001, 0.24, 0.25, respectively for 2, 4, 6 and 8 hours. (E) HHH014 and HHH014 *gyrB*^429N^. p = 0.02, 0.0008, 0.07, 0.0001, respectively for 2, 4, 6 and 8 hours. (F) FFF043 and FFF043 *gyrB*^429N^ (p value is not significant at all timepoints). (G) CCC033 and CCC033 *gyrB*^429N^. p = 0.31, 0.48, 0.27, 0.002, respectively for 2, 4, 6 and 8 hours. (H) DDD033 andDDD033 *gyrB*^429N^. p = 0.07, <0.0001, 0.85, 0.63, respectively for 2, 4, 6 and 8 hours. (I) HHH023 and HHH023 *gyrB*^429N^. p = 0.002, 0.42, 0.003, 0.01, respectively for 2, 4, 6 and 8 hours. n = 3, representative of three independent experiments. Error bars represent SDs between three biological replicates. Statistical significance was determined by unpaired two-sided Student’s *t-test* and are indicated *p ≤ 0.05, **p ≤ 0.01, ***p ≤ 0.001 and ****p ≤ 0.0001.

**Supplementary Figure 5:**
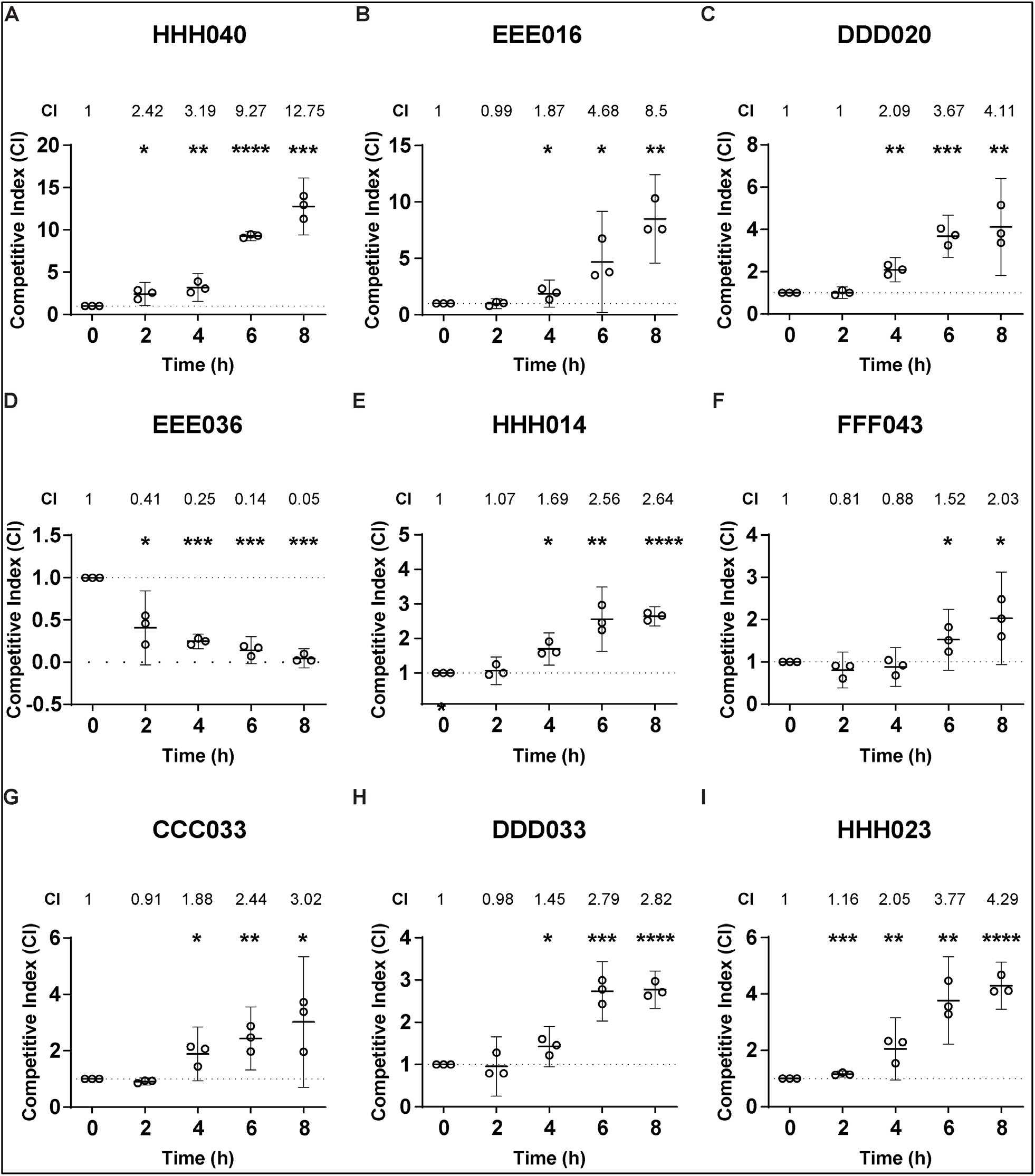
Relative fitness of the *gyrB*^D429N^ mutant in 9 clinical isolate back-grounds. Y-axes show the Competitive Index (CI) of each *gyrB*^D429N^ mutant relative to its parental strain during competitive growth *in vitro*. In all cases, the parental strain carried a kanamycin marker and was cocultured with its unmarked *gyrB*^D429N^ mutant. (A) HHH040: p = 0.01, 0.005, <0.0001, 0.0001, respectively for 2, 4, 6 and 8 hours. (B) EEE016: p = 0.9, 0.04, 0.02, 0.001, respectively for 2, 4, 6 and 8 hours. (C) DDD020: p = 0.92, 0.001, 0.0003, 0.004, respectively for 2, 4, 6 and 8 hours. (D) EEE036: p = 0.004, <0.0001, <0.0001, <0.0001, respectively for 2, 4, 6 and 8 hours. (E) HHH014: p = 0.51, 0.003, 0.002, <0.0001, respectively for 2, 4, 6 and 8 hours. (F) FFF043: p = 0.12, 0.32, 0.04, 0.02, respectively for 2, 4, 6 and 8 hours. (G) CCC033: p = 0.03, 0.02, 0.005, 0.02, respectively for 2, 4, 6 and 8 hours. (H) DDD033: p = 0.79, 0.02, 0.0004, <0.0001, respectively for 2, 4, 6 and 8 hours. (I) HHH023: p = 0.003, 0.01, 0.002, <0.0001, respectively for 2, 4, 6 and 8 hours. n = 3, representative of three independent experiments performed in absence of any antibiotic pressure. Error bars represent mean with 95% confidence interval. Statistically significant differences in competitive indices compared to time 0 were analyzed using an unpaired Student’s *t-test*, indicated *p ≤ 0.05, **p ≤ 0.01, ***p ≤ 0.001 and ****p ≤ 0.0001.

**Supplementary Table 1:**
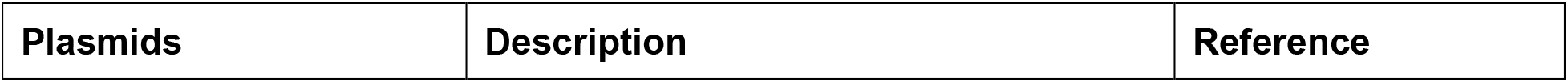
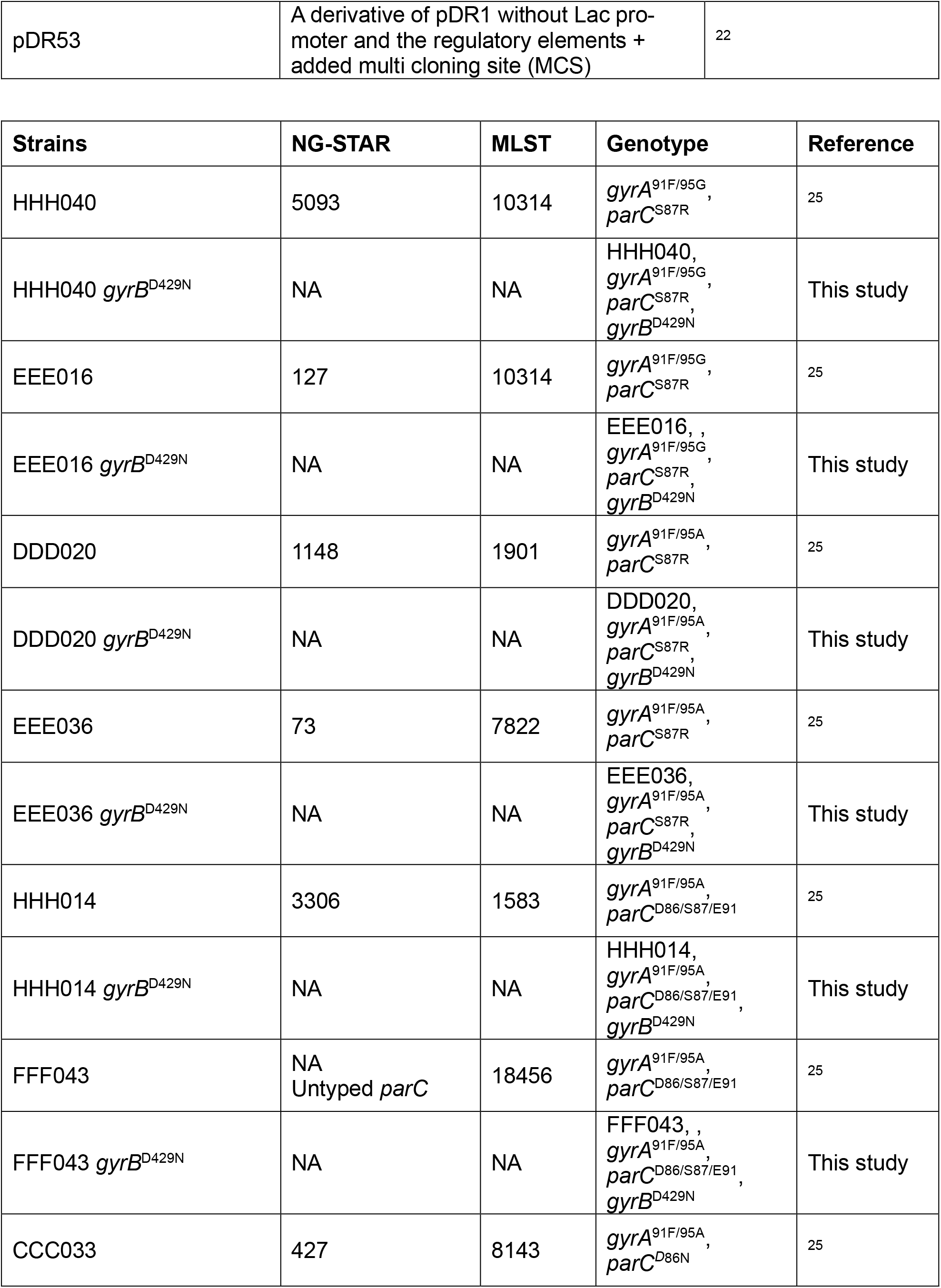
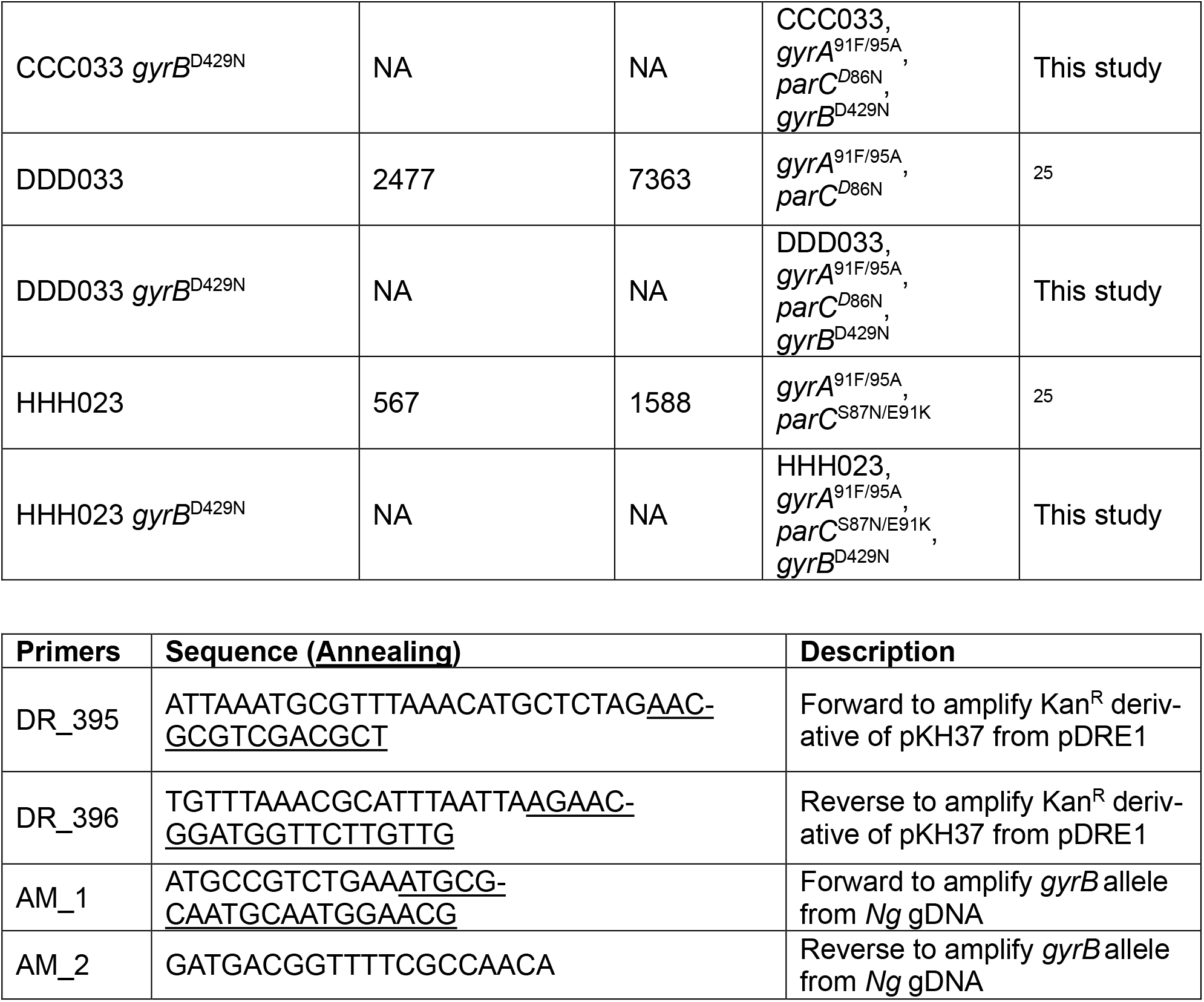
Plasmids, *N. gonorrhoeae* strains and primers used in this study.

**Supplementary Table 2:** Whole genome sequencing analysis of *gyrB*^D429N^ transformants.

| Mutant | Reference | Variants |
| --- | --- | --- |
| HHH040 *gyrB*<sup>D429N</sup> | HHH040 | *gyrB*<sup>D429N</sup>, one *opa* pentanucleotide repeat variation*, repeat variation in gene of unknown function homologous to NCCP11945 NGK_RS02285 |
| HHH014 *gyrB*<sup>D429N</sup> | HHH014 | *gyrB*<sup>D429N</sup>, changes in two *opa* genes*, repeat variation in site-specific DNA methyltransferase M.NgoAXII* |
| HHH023 *gyrB*<sup>D429N</sup> | HHH023 | *gyrB*<sup>D429N</sup>, single nucleotide deletion in intergenic region between IS110-family transposase and hypothetical protein homologous to NCCP11945 NGK_RS06655 |
| EEE016 *gyrB*<sup>D429N</sup> | EEE016 | *gyrB*<sup>D429N</sup>, *pilT* SNP, SNP in a transposase pseudogene, repeat variation in gene of unknown function homologous to NCCP11945 NGK_RS02285, small insertion in *mexA*, changes in one *opa* gene*, repeat variation in site-specific DNA methyltransferase M.NgoAXII* |
| FFF043 *gyrB*<sup>D429N</sup> | FFF043 | *gyrB*<sup>D429N</sup>, *pilE* SNP, repeat variation in site-specific DNA methyltransferase M.NgoAXII* |
| DDD020 *gyrB*<sup>D429N</sup> | DDD020 | *gyrB*<sup>D429N</sup>, no others |
| DDD033 *gyrB*<sup>D429N</sup> | DDD033 | *gyrB*<sup>D429N</sup>, one *opa* pentanucleotide repeat variation*, repeat variation in site-specific DNA methyltransferase M.NgoAXII* |
| CCC033 *gyrB*<sup>D429N</sup> | CCC033 | *gyrB*<sup>D429N</sup>, no others |
| EEE036 *gyrB*<sup>D429N</sup> | EEE036 | *gyrB*<sup>D429N</sup>, no others |

\* High frequency variation at *opa* and *modA* (encoding M.NgoXII) loci is known to occur via changes in repeat copy number<sup>49, 50</sup> and via intergenic recombination (for *opa*)<sup>51</sup>. Variation in *opa* and *modA* is known to influence adhesion (e.g., reviewed in<sup>52</sup>) and epigenetic programming<sup>50</sup> of *N. gonorrhoeae*, respectively. Neither *opa* nor *modA* phase variation in the *gyrB*<sup>D429N</sup> derivatives we studied here correlated with changes in resistance (Figure 1A) or fitness (Figure 2).

## References

1. Gandra, S. et al. Antimicrobial Resistance Surveillance in Low- and Middle-Income Countries: Progress and Challenges in Eight South Asian and Southeast Asian Countries. Clin Microbiol Rev 33 (2020).

2. Zhu, X., Xi, Y., Gong, X. & Chen, S. Ceftriaxone-Resistant Gonorrhea - China, 2022. MMWR Morb Mortal Wkly Rep 73, 255–259 (2024).

3. Laumen, J.G.E. et al. High Prevalence of Ceftriaxone-Resistant Neisseria gonorrhoeae in Hanoi, Vietnam, 2023-2024. J Infect Dis 232, e73–e77 (2025).

4. Sarah McLeod, P., Varalakshmi Elango, M., Esther Bettiol, MD PhD, Alison Luckey, M. (Open Forum Infectious Diseases, 2025).

5. Ross, J.D.C. et al. Oral gepotidacin for the treatment of uncomplicated urogenital gonorrhoea (EAGLE-1): a phase 3 randomised, open-label, non-inferiority, multicentre study. Lancet 405, 1608–1620 (2025).

6. Oviatt, A.A. et al. Mechanism of Action of Gepotidacin: Well-Balanced Dual-Targeting against Neisseria gonorrhoeae Gyrase and Topoisomerase IV in Cells and In Vitro. ACS Infect Dis (2025).

7. Alm, R.A. et al. Characterization of the novel DNA gyrase inhibitor AZD0914: low resistance potential and lack of cross-resistance in Neisseria gonorrhoeae. Antimicrob Agents Chemother 59, 1478–1486 (2015).

8. Belland, R.J., Morrison, S.G., Ison, C. & Huang, W.M. Neisseria gonorrhoeae acquires mutations in analogous regions of gyrA and parC in fluoroquinolone-resistant isolates. Mol Microbiol 14, 371–380 (1994).

9. Rubin, D.H., Mortimer, T.D. & Grad, Y.H. Neisseria gonorrhoeae diagnostic escape from a gyrA-based test for ciprofloxacin susceptibility and the effect on zoliflodacin resistance: a bacterial genetics and experimental evolution study. Lancet Microbe 4, e247–e254 (2023).

10. Foerster, S. et al. In vitro antimicrobial combination testing of and evolution of resistance to the first-in-class spiropyrimidinetrione zoliflodacin combined with six therapeutically relevant antimicrobials for Neisseria gonorrhoeae. J Antimicrob Chemother 74, 3521–3529 (2019).

11. Jacobsson, S. et al. Pharmacodynamic Evaluation of Zoliflodacin Treatment of Neisseria gonorrhoeae Strains With Amino Acid Substitutions in the Zoliflodacin Target GyrB Using a Dynamic Hollow Fiber Infection Model. Front Pharmacol 13, 874176 (2022).

12. Taylor, S.N. et al. Gepotidacin for the Treatment of Uncomplicated Urogenital Gonorrhea: A Phase 2, Randomized, Dose-Ranging, Single-Oral Dose Evaluation. Clin Infect Dis 67, 504–512 (2018).

13. Scangarella-Oman, N.E. et al. Dose selection for a phase III study evaluating gepotidacin (GSK2140944) in the treatment of uncomplicated urogenital gonorrhoea. Sex Transm Infect 99, 64–69 (2023).

14. Scangarella-Oman, N.E. et al. Microbiological Analysis from a Phase 2 Randomized Study in Adults Evaluating Single Oral Doses of Gepotidacin in the Treatment of Uncomplicated Urogenital Gonorrhea Caused by Neisseria gonorrhoeae. Antimicrob Agents Chemother 62 (2018).

15. Jacobsson, S., Golparian, D., Scangarella-Oman, N. & Unemo, M. In vitro activity of the novel triazaacenaphthylene gepotidacin (GSK2140944) against MDR Neisseria gonorrhoeae. J Antimicrob Chemother 73, 2072–2077 (2018).

16. Bradford, P.A., Miller, A.A., O’Donnell, J. & Mueller, J.P. Zoliflodacin: An Oral Spiropyrimidinetrione Antibiotic for the Treatment of Neisseria gonorrheae, Including Multi-Drug-Resistant Isolates. ACS Infect Dis 6, 1332–1345 (2020).

17. Lan, P.T. et al. The WHO Enhanced Gonococcal Antimicrobial Surveillance Programme (EGASP) identifies high levels of ceftriaxone resistance across Vietnam, 2023. Lancet Reg Health West Pac 48, 101125 (2024).

18. Ouk, V. et al. World Health Organization Enhanced Gonococcal Antimicrobial Surveillance Programme, Cambodia, 2023. Emerg Infect Dis 30, 1493–1495 (2024).

19. Golparian, D. et al. Genomic surveillance and antimicrobial resistance in Neisseria gonorrhoeae isolates in Bangkok, Thailand in 2018. J Antimicrob Chemother 77, 2171–2182 (2022).

20. Reichert, E. et al. Resistance-minimising strategies for introducing a novel antibiotic for gonorrhoea treatment: a mathematical modelling study. Lancet Microbe 4, e781–e789 (2023).

21. Kline, M.C., Roster, K.O., Helekal, D., Rumpler, E. & Grad, Y.H. Comparing Strategies to Introduce Two New Antibiotics for Gonorrhea: A Modeling Study. medRxiv (2025).

22. Helekal, D. et al. Quantifying the impact of antibiotic use and genetic determinants of resistance on bacterial lineage dynamics. bioRxiv (2025).

23. Rubin, D.H.F. et al. CanB is a metabolic mediator of antibiotic resistance in Neisseria gonorrhoeae. Nat Microbiol 8, 28–39 (2023).

24. Vegvari, C. et al. Using rapid point-of-care tests to inform antibiotic choice to mitigate drug resistance in gonorrhoea. Euro Surveill 25 (2020).

25. Bristow, C.C. et al. Whole-Genome Sequencing to Predict Antimicrobial Susceptibility Profiles in Neisseria gonorrhoeae. J Infect Dis 227, 917–925 (2023).

26. Lan, P.T. et al. Genomic analysis and antimicrobial resistance of Neisseria gonorrhoeae isolates from Vietnam in 2011 and 2015-16. J Antimicrob Chemother 75, 1432–1438 (2020).

27. Golparian, D. et al. Antimicrobial-resistant Neisseria gonorrhoeae in Europe in 2020 compared with in 2013 and 2018: a retrospective genomic surveillance study. Lancet Microbe 5, e478–e488 (2024).

28. Aligning sequence reads, clone sequences and assembly contigs with BWA-MEM. (2013).

29. Walker, B.J. et al. Pilon: an integrated tool for comprehensive microbial variant detection and genome assembly improvement. PLoS One 9, e112963 (2014).

30. Li, H. et al. The Sequence Alignment/Map format and SAMtools. Bioinformatics 25, 2078–2079 (2009).

31. Croucher, N.J. et al. Rapid phylogenetic analysis of large samples of recombinant bacterial whole genome sequences using Gubbins. Nucleic Acids Res 43, e15 (2015).

32. Minh, B.Q. et al. IQ-TREE 2: New Models and Efficient Methods for Phylogenetic Inference in the Genomic Era. Mol Biol Evol 37, 1530–1534 (2020).

33. Kalyaanamoorthy, S., Minh, B.Q., Wong, T.K.F., von Haeseler, A. & Jermiin, L.S. ModelFinder: fast model selection for accurate phylogenetic estimates. Nat Methods 14, 587–589 (2017).

34. Letunic, I. & Bork, P. Interactive Tree of Life (iTOL) v6: recent updates to the phylogenetic tree display and annotation tool. Nucleic Acids Res 52, W78–W82 (2024).

35. Mirdita, M. et al. ColabFold: making protein folding accessible to all. Nat Methods 19, 679–682 (2022).

36. Kozakov, D. et al. The ClusPro web server for protein-protein docking. Nat Protoc 12, 255–278 (2017).

37. Pettersen, E.F. et al. UCSF ChimeraX: Structure visualization for researchers, educators, and developers. Protein Sci 30, 70–82 (2021).

38. Kellogg, D.S., Jr., Peacock, W.L., Jr., Deacon, W.E., Brown, L. & Pirkle, D.I. Neisseria Gonorrhoeae. I. Virulence Genetically Linked to Clonal Variation. J Bacteriol 85, 1274–1279 (1963).

39. Dillard, J.P. Genetic Manipulation of Neisseria gonorrhoeae. Curr Protoc Microbiol Chapter 4, Unit4A 2 (2011).

40. Wick, R.R., Howden, B.P. & Stinear, T.P. Autocycler: long-read consensus assembly for bacterial genomes. Bioinformatics 41 (2025).

41. Koren, S. et al. Canu: scalable and accurate long-read assembly via adaptive k-mer weighting and repeat separation. Genome Res 27, 722–736 (2017).

42. Kolmogorov, M., Yuan, J., Lin, Y. & Pevzner, P.A. Assembly of long, error-prone reads using repeat graphs. Nat Biotechnol 37, 540–546 (2019).

43. Li, H. Minimap and miniasm: fast mapping and de novo assembly for noisy long sequences. Bioinformatics 32, 2103–2110 (2016).

44. Chen, Y. et al. Efficient assembly of nanopore reads via highly accurate and intact error correction. Nat Commun 12, 60 (2021).

45. Hu, J. et al. NextDenovo: an efficient error correction and accurate assembly tool for noisy long reads. Genome Biol 25, 107 (2024).

46. Vaser, R. & Sikic, M. Time- and memory-efficient genome assembly with Raven. Nat Comput Sci 1, 332–336 (2021).

47. Li, H. Minimap2: pairwise alignment for nucleotide sequences. Bioinformatics 34, 3094–3100 (2018).

48. Seemann, T. Prokka: rapid prokaryotic genome annotation. Bioinformatics 30, 2068–2069 (2014).

49. Stern, A., Brown, M., Nickel, P. & Meyer, T.F. Opacity genes in Neisseria gonorrhoeae: control of phase and antigenic variation. Cell 47, 61–71 (1986).

50. Srikhanta, Y.N. et al. Phasevarions mediate random switching of gene expression in pathogenic Neisseria. PLoS Pathog 5, e1000400 (2009).

51. Connell, T.D. et al. Recombination among protein II genes of Neisseria gonorrhoeae generates new coding sequences and increases structural variability in the protein II family. Mol Microbiol 2, 227–236 (1988).

52. Walker, E., van Niekerk, S., Hanning, K., Kelton, W. & Hicks, J. Mechanisms of host manipulation by Neisseria gonorrhoeae. Front Microbiol 14, 1119834 (2023).

